# AFsample2: Predicting multiple conformations and ensembles with AlphaFold2

**DOI:** 10.1101/2024.05.28.596195

**Authors:** Yogesh Kalakoti, Björn Wallner

**Affiliations:** Division of Bioinformatics, Department of Physics, Chemistry and Biology, Linköping University, 581 83 Linköping, Sweden

**Keywords:** Sampling, Alphafold, Conformations, Ensembles, Diversity

## Abstract

Understanding protein dynamics and conformational states carries profound scientific and practical implications for several areas of research, ranging from a general understanding of biological processes at the molecular level to a detailed understanding of disease mechanisms, which in turn can open up new avenues in drug development. Multiple solutions have been recently developed to widen the conformational landscape of predictions made by Alphafold2 (AF2). Here, we introduce AFsample2, a method employing random MSA column masking to reduce the influence of co-evolutionary signals to enhance the structural diversity of models generated by the AF2 neural network. AFsample2 improves the prediction of alternative states for a broad range of proteins, yielding high-quality end states and diverse conformational ensembles. In the data set of open-closed conformations (OC23), alternate state models improved in 17 out of 23 cases without compromising the generation of the preferred state. Consistent results were observed in 16 membrane protein transporters, with improvements in 12 out of 16 targets. TM-score improvements to experimental end states were substantial, sometimes exceeding 50%, elevating mediocre scores from 0.58 to nearly perfect 0.98. Furthermore, AFsample2 increased the diversity of intermediate conformations by 70% compared to the standard AF2 system, producing highly confident models, that could potentially be on-path between the two states. In addition, we also propose a way of selecting the end-states in generated model ensembles. These solutions could potentially enhance the generation and identification of alternative protein conformations, thereby providing a more comprehensive understanding of protein function and dynamics. Future work will focus on validating the accuracy of these intermediate conformations and exploring their relevance to functional transitions in proteins.

## Introduction

Proteins are the workhorses of life, serving as the building blocks of cells and playing crucial roles in almost every biological process. They exhibit a wide range of functions, including catalyzing biochemical reactions, providing structural reinforcement, and even acting as conduits in intracellular communication (1). Proteins adopt intricate threedimensional configurations, often existing within structural ensembles that exhibit various states, collective movements, and dynamic fluctuations, all essential for executing their function (2, 3). Processes such as folding, signal transduction, enzyme catalysis, and molecular recognition are all driven by the type and extent of structural dynamics associated with the protein system. Conventional experimental structural biology methods such as X-ray crystallography and cryogenic electron microscopy (cryo-EM) can provide a few highly accurate snapshots of the overall conformational ensemble of the protein system (4–6). However, these snapshots only represent a fraction of possible states and have to be supplemented by molecular dynamics or other similar solutions to infer molecular mechanisms. Furthermore, computational costs related to MD at biologically relevant timescales are not viable in practice. Other experimental methods such as Nuclear Magnetic Resonance (NMR) could potentially profile the dynamic nature of the protein molecule, but are limited by scale (7).

Recent advancements in *in-silico* protein structure determination have largely been an outcome of intelligent data processing and generative artificial intelligence (AI). Methods like AlphaFold2 (8) (AF2) and RosettaFold (9) have demonstrated exceptional levels of success in determining accurate protein structure from evolutionary sequence information provided as Multiple Sequence Alignments (MSAs). However, the default versions of these workflows are trained to estimate a single high-confident model of the structure of a protein. This is a limitation since the entire conformational landscape has to be considered in order to get insights into the mechanistic basis of protein function. Therefore, an ideal sequence-to-structure prediction system should have the ability to model the entire conformational ensemble for a given protein, identify states, and trace physically viable paths in estimated ensembles. Our recently developed AFsample method (10) captured different conformations of multimeric proteins by increasing the sampling rate and introducing noise by enabling dropout layers at inference. The method achieved state-of-the-art performance and was one of the top-ranked at CASP15 (11) (2022). Additional strategies have also been proposed to induce conformational diversity in AF2 predictions by subsampling the MSA, using shallow MSAs (12), *in-silico* alanine scan as in SPEACH_AF (13), or clustering the MSA as in AFcluster (14). All of these methods work by effectively reducing the information to AF2 to allow the system to explore alternative solutions.

In this work, we present AFsample2, which employs random MSA column masking to diminish the constraints exerted by co-evolutionary signals in MSA. Thereby increasing the structural heterogeneity of models generated with the AF2 neural network. AFsample2 was able to improve the prediction of alternative states for a wide range of proteins.

The improvement was quantified based on the ability of the inference system to generate high-quality end states and diverse conformational ensembles. The models for in particular the alternate state is improved for most of the cases (17/23) in the open-closed data set (OC23) without sacrificing performance for the preferred state. The performance is maintain on an additional set of 16 membrane protein transporters, with the alternate state improved for 12/16 targets. The improvement as measured by TM-score to experimental end states is sometimes massive with improvements over 50%, basically going from mediocre TM-scores of 0.58 to almost perfect 0.98. However, the improvements are not only in the end-states but AFsample2 also improves the diversity by generating 70% more conformations in-between the end-states, when compared to the the vanilla AF2 system. While it remains to be demonstrated whether these intermediate conformations are accurate on-path representations between states, they are highly confident models generated by the AF2 inference system.

Furthermore, a novel strategy to identify conformational states from a pool of generated models without the aid of any experimental reference structures was also developed. In summary, significant methodological improvements have been presented in this study that enhance the capability of MSA-based generative models in capturing the conformational landscape of a given protein system.

## Results

### Method Development

The primary objective of this study was to improve the sampling of conformational states by introducing more noise than simply turning on the dropout layers at inference. In AFsample2, the noise is introduced by randomly masking columns in the MSA, with the rationale to break covariance constraints in the MSA, see Fig 1a. By breaking covariance signals, the inference system is allowed to explore and arrive at different solutions for the given protein, ultimately increasing the diversity of the generated protein ensemble. A similar strategy for introducing noise to MSAs has previously been attempted with SPEACH_AF (13), where a sliding window of alanines was introduced at specific columns in the MSA to break interacting residues. Although effective, this strategy was dependent on *in-silico* mutagenesis of the MSA, requiring prior knowledge of the interacting residues. In their implementation, these residues were based on either prior structural informa-tion or/and contacts in generated models. AFsample2 does not have this limitation and provides a general solution even for cases where such information is unavailable.

**Fig. 1.**
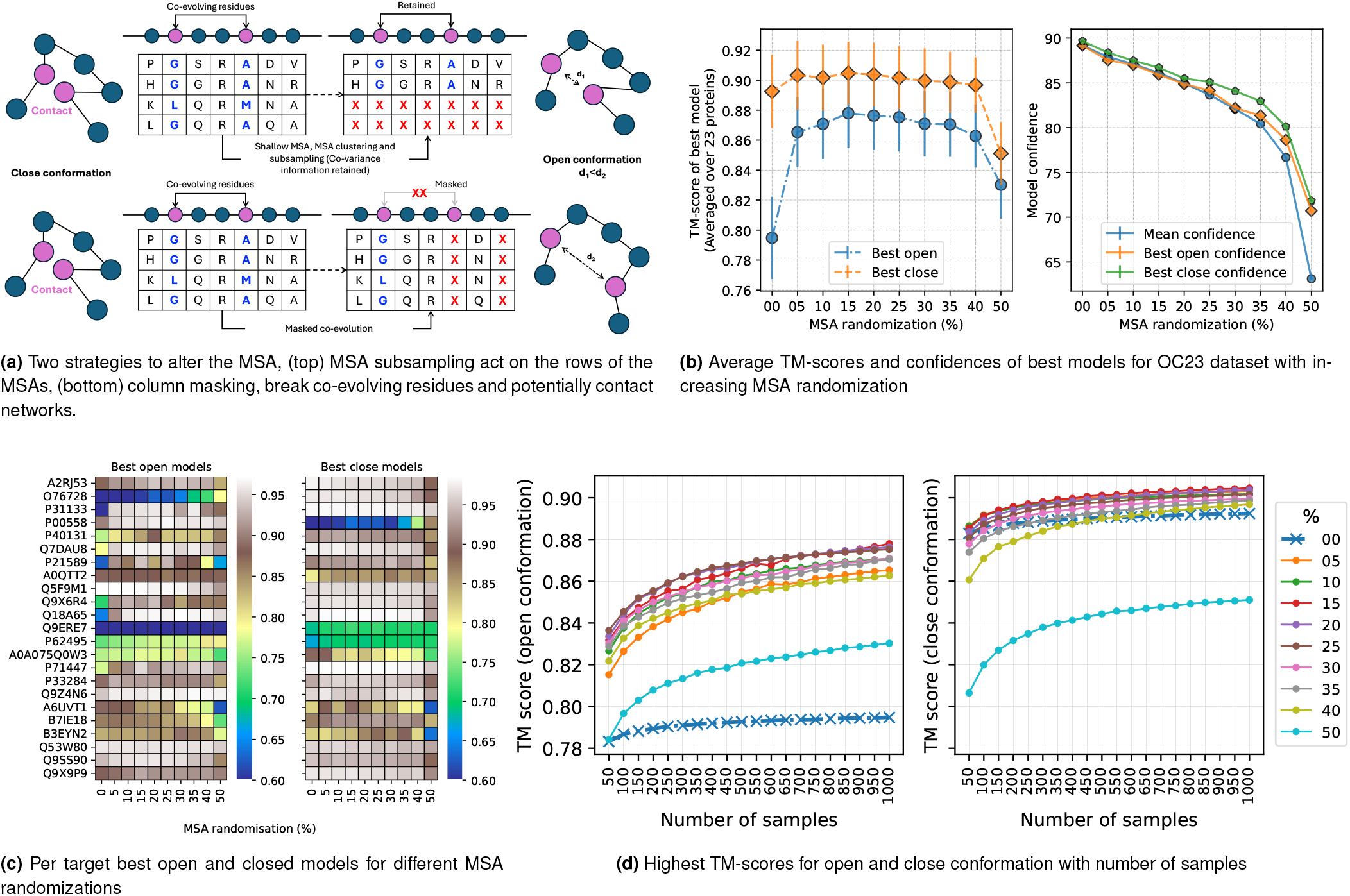
Overall summary and analysis of MSA randomization strategy in AFsample2. (a) A general outline of the modifications to the MSA for the AFsample2 pipeline (bottom). Traditional methods retain the information on co-variation, which in turn constrains the inference system to generate structures. AFsample2, on the other hand, remove those constraints by masking columns in the MSA to partially remove this co-variance information, leading to the generation of alternate conformations. (b) Effectiveness of the randomization strategy in terms of generating high-quality models and aggregate confidence for both open and closed states. The results indicate 15% randomization to have the highest TM-scores in the OC23 dataset. (c) Optimal level of randomization on a per target protein. (d) Sampling more models increases the chances of generating better models, and is significantly more potent with the proposed randomizations.

#### Effect of MSA masking on generated models

The amount of MSA masking, i.e., the fraction of randomized positions, was observed to be the most important factor in the ability of the inference system to generate alternate conformational states. It was observed that increasing the MSA masking increased the chances of generating end-state conformations for a given protein. This trend is summarized in Fig 1b, where MSA masking generates significantly better models compared to no masking (0%) for the alternate state (open in these cases) across a set of diverse proteins with well-defined open and closed conformations (the OC23 set, see Methods). The aggregate TM-score for the best alternate (open) conformation increases from 0.795 for no masking to 0.878 with 15% masking while showing a marginal improvement from 0.89 to 0.90 for the closed conformation. Beyond 30% masking, performance drops first for the open conformations and subsequently for the closed conformations.

In addition, it has previously been reported that the model confidence of AF2 predictions deteriorates with increased sub-sampling (15). A similar trend was also observed here, where the mean confidence gradually decreased with increasing MSA masking (Fig. 1b). The decrease is linear from 0% to 35% with a 2% drop in confidence for every 5 percentage points of masking, followed by a rapid drop in model confidence beyond 35% masking. Since MSA masking essentially removes information, this trend is expected. However, it is important to realize that the decrease in model confidence up to 20% masking is not coupled to lower quality models. It is most likely an effect that the mask itself renders more uncertainty in the prediction, which in turn results in lower model confidence. Overall, 15% randomization seems to perform marginally better than other settings. However, by analyzing the per target performance (Fig. 1c), it can be seen that different levels of masking yield the best performance for different target proteins, e.g., 20% masking generates the best model for P40131, while 5% masking is optimal for P71147. The best TM-scores for each protein at various masking levels can be visualized in Fig. S1. Even though the exact magnitude of masking might differ between targets, it is true that in all cases, masking is always better than no masking for the same level of sampling. For comparative analysis and simplicity, AFsample2 using 15% masking was used for the downstream analysis.

#### The importance of sampling

It has been previously established that increased sampling improves the chances of generating alternative conformations (10, 16). But the question is, how much sampling is enough? To answer this question, we estimated the best TM-score for the open and closed states, respectively, as the number of samples increased and for different levels of masking 0-50%, see Fig. 1d. Indeed, generating more sampling increases the chances of generating higher-quality models for all levels of masking. The improvement is larger for predicting the open conformation, re-flecting the fact that AF2 has a preference for predicting the closed conformation in this case, leaving more room for improvement of the open conformation. The increasing trend is most pronounced for fewer samples, reflecting the switch from no sampling to actual sampling, but it is still increasing even up to 1000 samples, indicating that sampling more is always better. However, considering the trade-off between speed and performance, 1000 samples at 15% masking is a reasonable default.

#### Overview of the AFsample2

Given a protein sequence, AFsample2 follows a four-step process to generate diverse protein structures using a modified version of the AF2 inference system. It starts by (i) querying sequence databases to generate multiple sequence alignments (MSAs), (ii) Randomly masking MSA columns with a pre-defined probability (e.g. 15%), (iii) running inference on a uniquely masked MSA for each model and lastly, (iv) depending on the availability of reference states, identifying state representatives with clustering, confidence and extremity selection, followed by ensemble analysis. A schematic representation of the workflow is summarized in Fig. 2.

**Fig. 2.**
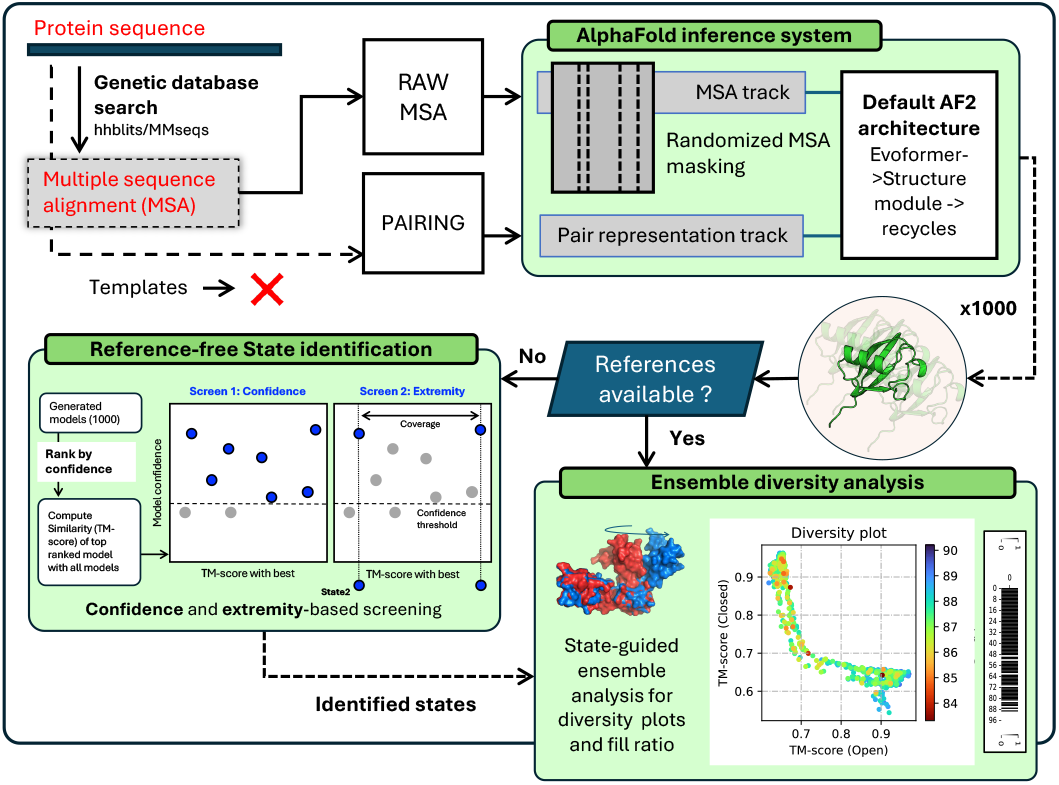
Overall workflow of the AFsample2 pipeline. It starts by generating MSAs for a given protein sequence. This is followed by randomized MSA masking in a way such that a unique MSA profile is fed into the system at every instance of the inference run. The generated ensemble is either passed to the diversity analysis workflow or the state-identification workflow, depending on the availability of reference states.

### Comparing AFsample2 to AFvanilla, AFdropout and Afcluster

The performance of AFsample2 was compared to standard AF2 (AFvanilla), standard AF2 with dropout (AFdropout), and AFcluster (14) by generating 1000 models for each protein in the OC23 dataset. Fig. 3a shows the distribution of TM-scores for the open and closed states for the models generated by the four methods. In terms of sampling different states, the performance of AFvanilla and AFdropout is almost identical; both generally have fairly narrow distributions centered on high TM-score for the closed state. In contrast, the distributions from AFsample2 cover a wider range, and importantly, the distributions for the open state cover the high TM-score region. This is unlike AFcluster, where even though the distributions are wider, a significant fraction of the ensemble can be attributed to bad-quality models (low TM-score for both states).

**Fig. 3.**
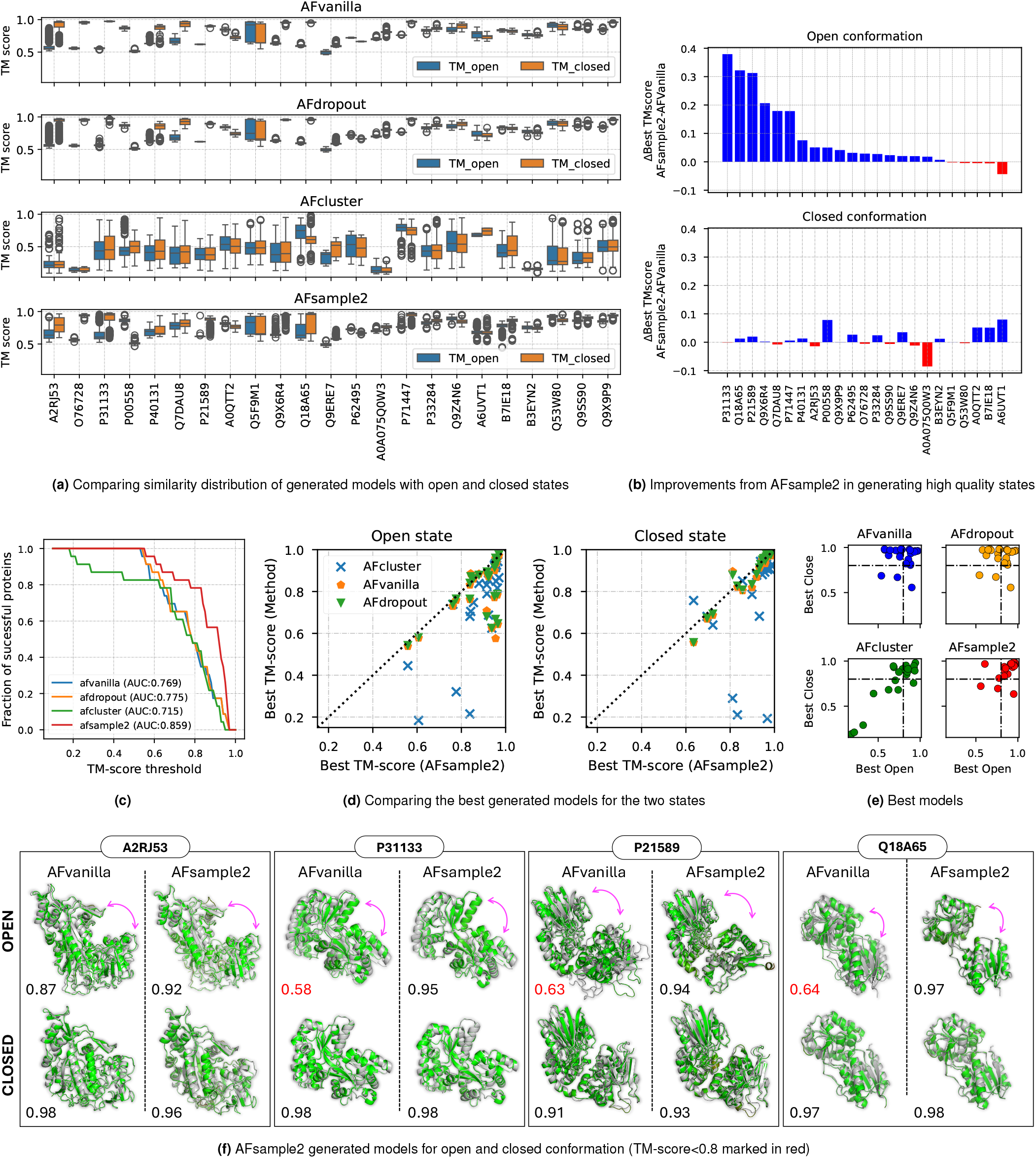
AFsample2 demonstrates a greater ability to generate good-quality open and closed conformations than baseline methods. (a) Distribution of TM-scores with reference open and closed structures with generated models for all targets in the OC23 dataset. (b) Differences between TM-scores of best open and closed models generated by AFsample2 and AFvanilla show improvements induced by MSA-masking (blue), while relatively deficient models were observed in a handful of cases (red). (c) Comparing TM-scores of best models generated for open and closed states by all methods under consideration. (d) Fraction of successful targets compared among methods at different levels of TM-score thresholds. Success: TM-score > threshold (for both states) (d) Comparing best generated open and closed models among methods. (e) Distribution of best models among methods. (e) Examples of best open and closed models that were generated by AFvanilla and AFsample2 with the same amount of sampling.

By analyzing the difference between the TM-score for the best open and closed models for AFsample2 to AFvanilla, see Fig. 3b. It is clear that for most of the targets, AFsample2 is able to generate better conformations of the open state without the best-closed conformations getting much worse. Only for a single case does it get worse than -0.1 TM-score units. This is further confirmed with Fig. 3c, where the level of success, measured by having both states with TM-score*>*threshold, is compared for the four methods at different thresholds. AFsample2 generates models of both states with TM-score*>*0.8 for 78.3% of the targets, while AFvanilla, AFdropout, and AFcluster only do it for 47.8% of the targets. This is further visualized in Fig. 3d,e, where the best open and closed models for each target are shown for AFvanilla, AFdropout, AFcluster, and AFsample2. While AFvanilla and AFdropout show a similar preference towards the closed conformation, AFcluster seems to fail (TM-score to both states<0.5) on a few cases while successfully generating both conformations for other cases. On the other hand, AFsample2 is able to successfully generate high-quality models of both states without failing for even a single target in the OC23 dataset.

Furthermore, Fig. 3e exemplifies a few successful AFsample2 predictions (green) superimposed on the open and closed experimental references (gray). It is quite evident that AFsample2 is able to generate high-accuracy models of both states (TM-score between generated model and reference *>* 0.9). This is a significant improvement over AFvanilla, where there seems to be a clear preference towards the closed state in the depicted examples. This trend is mostly consistent for all targets in the OC23 dataset, and a summary of best-generated states is compiled in Table 1.

**Table 1.** Summary of the results comparing the similarity of best open and closed models generated by AFvanilla and AFsample2.The first set is the group of proteins that did relatively high mean similarity (TM-score) with the experimental open and closed structures, and the second set consists of the proteins that could not generate good quality structures with AFvanilla. The improvements observed with AFsample2 are also summarised for both cases. Positive values in the Δ column indicate an increase in performance and vice versa.

| Target | BaselineTM<br>between states | Similarity to open conformation |  |  | Similarity to closed conformation |  |  |
| --- | --- | --- | --- | --- | --- | --- | --- |
| | | AFvanilla | AFsample2 | $\Delta$ | AFvanilla | AFsample2 | $\Delta$ |
| A2RJ53 | 0.507 | 0.872 | 0.939 | 0.067 | 0.978 | 0.965 | -0.013 |
| O76728 | 0.523 | 0.578 | 0.627 | 0.049 | 0.973 | 0.967 | -0.007 |
| P31133 | 0.565 | 0.576 | 0.959 | 0.383 | 0.981 | 0.979 | -0.003 |
| P00558 | 0.598 | 0.901 | 0.962 | 0.061 | 0.557 | 0.623 | 0.066 |
| P40131 | 0.612 | 0.763 | 0.881 | 0.119 | 0.924 | 0.923 | -0.001 |
| Q7DAU8 | 0.617 | 0.784 | 0.963 | 0.179 | 0.977 | 0.972 | -0.005 |
| P21589 | 0.621 | 0.625 | 0.835 | 0.210 | 0.912 | 0.93 | 0.017 |
| A0QTT2 | 0.633 | 0.884 | 0.878 | -0.006 | 0.804 | 0.851 | 0.047 |
| Q5F9M1 | 0.636 | 0.972 | 0.964 | -0.008 | 0.963 | 0.967 | 0.004 |
| Q9X6R4 | 0.648 | 0.708 | 0.935 | 0.227 | 0.968 | 0.967 | -0.001 |
| Q18A65 | 0.656 | 0.643 | 0.966 | 0.323 | 0.971 | 0.984 | 0.013 |
| Q9ERE7 | 0.664 | 0.538 | 0.548 | 0.010 | 0.687 | 0.709 | 0.022 |
| P62495 | 0.708 | 0.730 | 0.770 | 0.040 | 0.669 | 0.692 | 0.023 |
| A0A075Q0W3 | 0.720 | 0.759 | 0.776 | 0.017 | 0.896 | 0.831 | -0.065 |
| P71447 | 0.748 | 0.771 | 0.911 | 0.140 | 0.980 | 0.984 | 0.004 |
| P33284 | 0.749 | 0.888 | 0.918 | 0.030 | 0.934 | 0.96 | 0.026 |
| Q9Z4N6 | 0.749 | 0.939 | 0.964 | 0.025 | 0.962 | 0.957 | -0.005 |
| A6UVT1 | 0.754 | 0.886 | 0.838 | -0.048 | 0.819 | 0.861 | 0.042 |
| B7IE18 | 0.755 | 0.86 | 0.854 | -0.006 | 0.868 | 0.926 | 0.058 |
| B3EYN2 | 0.766 | 0.832 | 0.847 | 0.015 | 0.821 | 0.863 | 0.042 |
| Q53W80 | 0.781 | 0.958 | 0.956 | -0.003 | 0.960 | 0.959 | -0.0 |
| Q9SS90 | 0.824 | 0.934 | 0.958 | 0.024 | 0.951 | 0.947 | -0.005 |
| Q9X9P9 | 0.837 | 0.881 | 0.909 | 0.027 | 0.971 | 0.968 | -0.004 |
| Mean | 0.681 | 0.795 | 0.876 | 0.082 | 0.892 | 0.904 | 0.011 |

### AFsample2 generates diverse protein ensembles

The effectiveness of AFsample2 in generating open and closed conformations has been presented above. However, the analysis only considered the best open and closed models generated, completely ignoring the ensemble of models between the two states. This ensemble of models might capture relevant micro-states that are on-path between the two states, or they might simply be unsuccessful attempts to model the different states. In order to capture the presence or absence of diversity, each model in the generated ensemble was compared with reference open and closed states using TM-align and visualized on a scatter plot (*Diversity* plot). For instance, while considering the example of target P31133 (Putrescine-binding periplasmic protein PotF) in Fig. 4a, it is quite ev-ident that all methods successfully generate the closed con-formation (top left region in the diversity plot), however, AF-sample2 is able to generate both states, as well as all the prospective states between the two conformations (open and closed). On closer inspection of Fig. 4a, AFdropout seems to tend towards the alternate state but fails, possibly due to a sharp decline in model confidence. On the other hand, while the aggregate confidence of AFsample2 models is marginally lower, it is high enough to sample the complete entire space between the two states. Diversity plots for all targets in the OC23 dataset are compiled in Fig. S2.

**Fig. 4.**
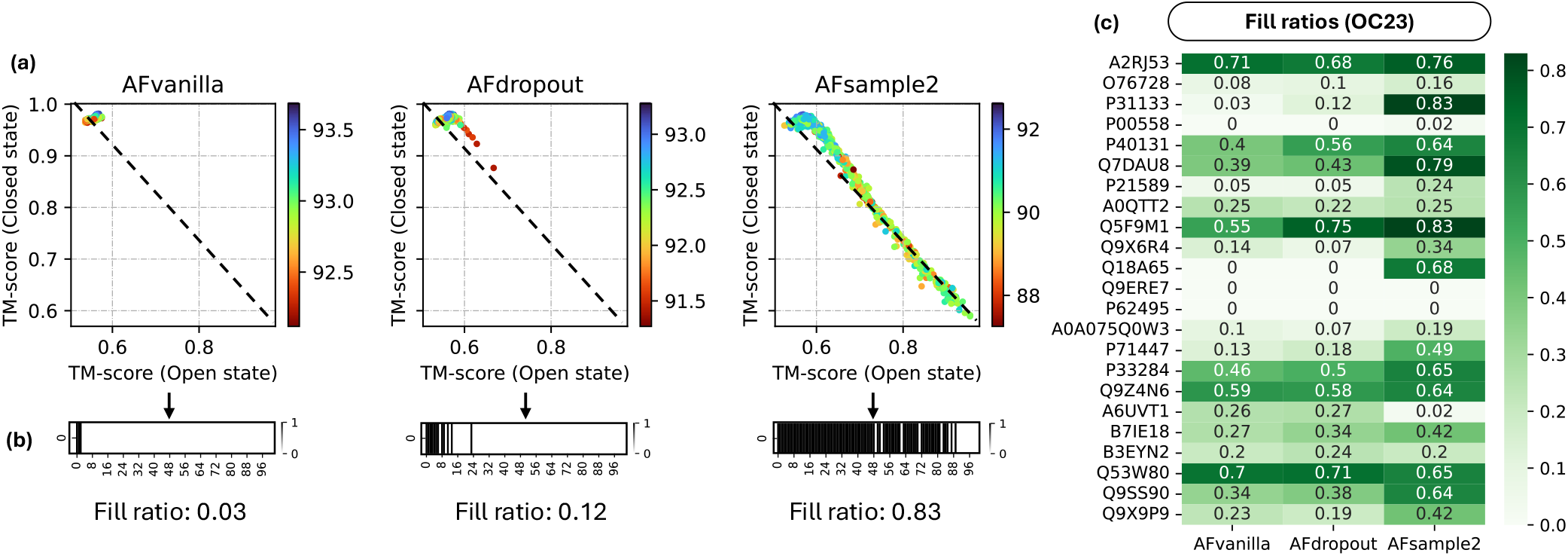
Analysing diversity of generated ensembles. (a) Diversity plot showing similarity of generated models with open and closed reference structures for AFvanilla, AFdropout and AFsample2 (Colorbar: model confidence). (b) Visualizing *fill-ratio*, a measure to quantify the fraction of diagonal covered by models in the diversity plot. (c) Heat-map summarizing *fill-ratios* for all targets in the OC23 dataset when compared between methods.

Although the presented example and visual cues from diversity plots (Fig. 4a) indicate the effectiveness of AFsample2 in generating diverse models, a measure that captures the extent of diversity is required for a valid comparison. In order to do so, a *fill-ratio* measure was developed. In short, the *fill-ratio* breaks down the path from open to closed state into 100 bins and calculates the percentage of bins that are populated by at least one model (as illustrated in Fig. 4b). Fig. 4c summarizes the *fill-ratios* for all targets in the OC23 dataset for AFvanilla, AFdropout, and AFsample2. Furthermore, diversity plots compiled in Fig. **??** indicate that while closed conformations are still the majority of the model population, MSA masking induces the inference system to have a higher probability of sampling alternate states at a given level of sampling.

### Fluctuations in predicted protein models agree with experimental data

As discussed earlier, AFsample2 induces randomness in the AF2 system as an attempt to diversify model predictions. However, it is not given that the diversity in the ensemble of models would actually agree with the experimentally observed fluctuations. To investigate this relationship, the Root Mean Squared Fluctuation (RMSF) of C*α* coordinates for generated models superimposed on the model with the highest model confidence was calculated for AFvanilla, AFdropout, and AFsample2. The RMSF profiles were compared to the actual per-residue distance between the opened and closed states (ΔC*α*), exemplified in Fig. 5a-d for target Q9X6R4. The correlation between the per-residue fluctuation and the RMSF profile for the example is 0.90, 0.69, and 0.96 for AFvanilla, AFdropout, and AFsample2, respectively. Overall, the correlation coefficients between the RMSF profile and ΔC*α* for all targets in the OC23 dataset are summarized in Fig. 5e. AFsample2 has a higher correlation for 19 out of 23 targets compared to either AFvanilla or AFdropout and correlates well (R*>*0.8) for 14 targets compared to only eight and seven for AFvanilla and AFdropout, respectively. While correlation is important, it does not take the magnitude of the movements into account, illustrated in Fig. 5b-d, where AFvanilla and AFsample both correlate well, but where AFsample2 is much better calibrated since the magnitude of fluctuations agrees much better.

**Fig. 5.**
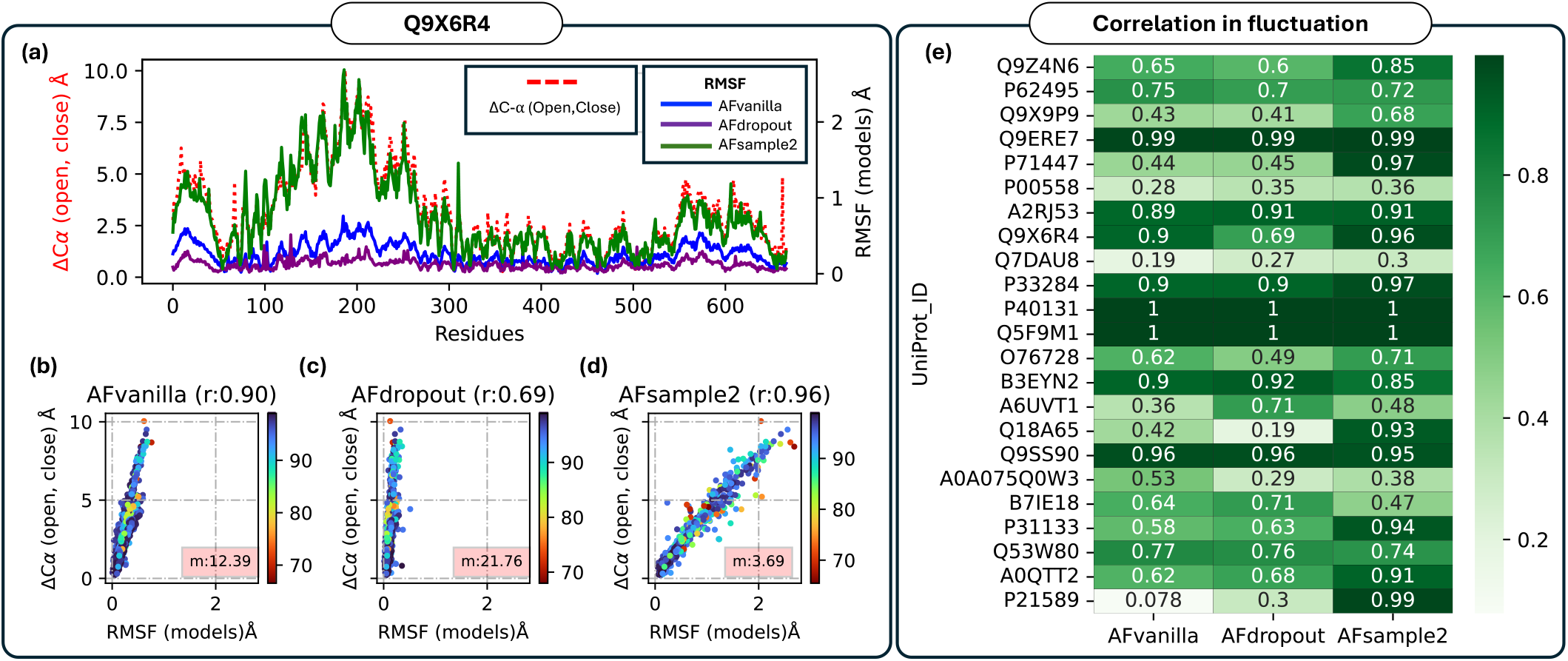
Comparing ensemble fluctuations with reference states. (a) A consolidated analysis of per-residue fluctuations observed in the model ensemble for Q9X6R4 and its correlation with C-*α* distances between experimental states, compared across AFvanilla, AFdropout and AFsample2 (b) Correlation statistics for all proteins in the OC23 dataset

If we assume that the fluctuations are normally distributed with RMSFs as standard deviations and that the ΔC*α* is, for instance, the 99.9th percentile of the distribution, the Z-score for ΔC*α* would be 3.29, i.e., Δ*Cα* = 3.29*RMSF* . This means that if you have the RMSF value, you should multiply it by 3.29 to get the ΔC*α*. It is not given that the ΔC*α* should be at the 99.9th percentile, but since the movement here is between an open and closed state, it is fair to assume that the residue distances between these states are among the largest and a Z-score of ≈3 should be expected if correct ΔC*α* have been generated from a normal distribution with RMSF as the standard deviation. If the Z-scores are much larger than 3, it would mean that the fluctuations are smaller than expected. The slope (m-value) of a linear fit between RMSF and ΔC*α* will give the Z-score and allow for a comparison of the magnitude of the movement. In Fig. 5b-d, the Z-score is 12.39, 21.76, and 3.69 for AFvanilla, AFdropout, and AFsample2, respectively, which indicate that the movement sampled by AFsample2 corresponds to would be expected, while the movement of AFvanilla and AFdropout is actually much smaller than expected. In fact, among all the 23 targets in the OC23 set, AFsample2 has an m-value between 2-5 for 16/23 targets, compared to 8/23 and 7/23 for AFvanilla and AFdropout, respectively, see Fig S5.

This observation is particularly interesting because the inference system relies solely on randomized MSAs without any additional information about the biological tendencies of fluctuations at the residue level. Still, the disruptions in co-variance signals caused by MSA masking seem to encourage backbone fluctuations, and the scoring system is able to dis-criminate between relevant and non-relevant fluctuations.

### Identify protein states without ground truths

Given a pool of generated models, it is non-trivial to correctly identify conformational states without the availability of reference structures. Current strategies rely on experimental reference structures to determine whether a given sampling method can generate alternate conformations. However, in the absence of the experimental reference structures, there is currently no standard way of determining valid states from the generated model ensemble. AFsample2 introduces a novel strategy to identify conformational states given a pool of models without the aid of experimentally solved reference structures. The method relies on the tendency of the AF2 inference system to favor a particular state by default. Assuming the AF2 inference system ranks one state higher in terms of confidence and the AFsample2 system promotes diversity, models in the alternative state are expected to differ significantly from the best model in terms of structural similarity (TM-score). Consequently, the alternative state should appear as a high-confidence model that is structurally distinct from the best model.

In short, the alternate state identification algorithm in AFsample2 follows a three-step process, starting with (i) calculating the similarity to the best model for all models in the ensemble, (ii) *Confidence screening* - for filtering models below a certain threshold, and (iii) *Extremity selection* - to identifying the model that is furthest from the most confident model. A visual representation of the idea of selected high-scoring models structurally dissimilar from the best model is depicted in Fig 6a.

**Fig. 6.**
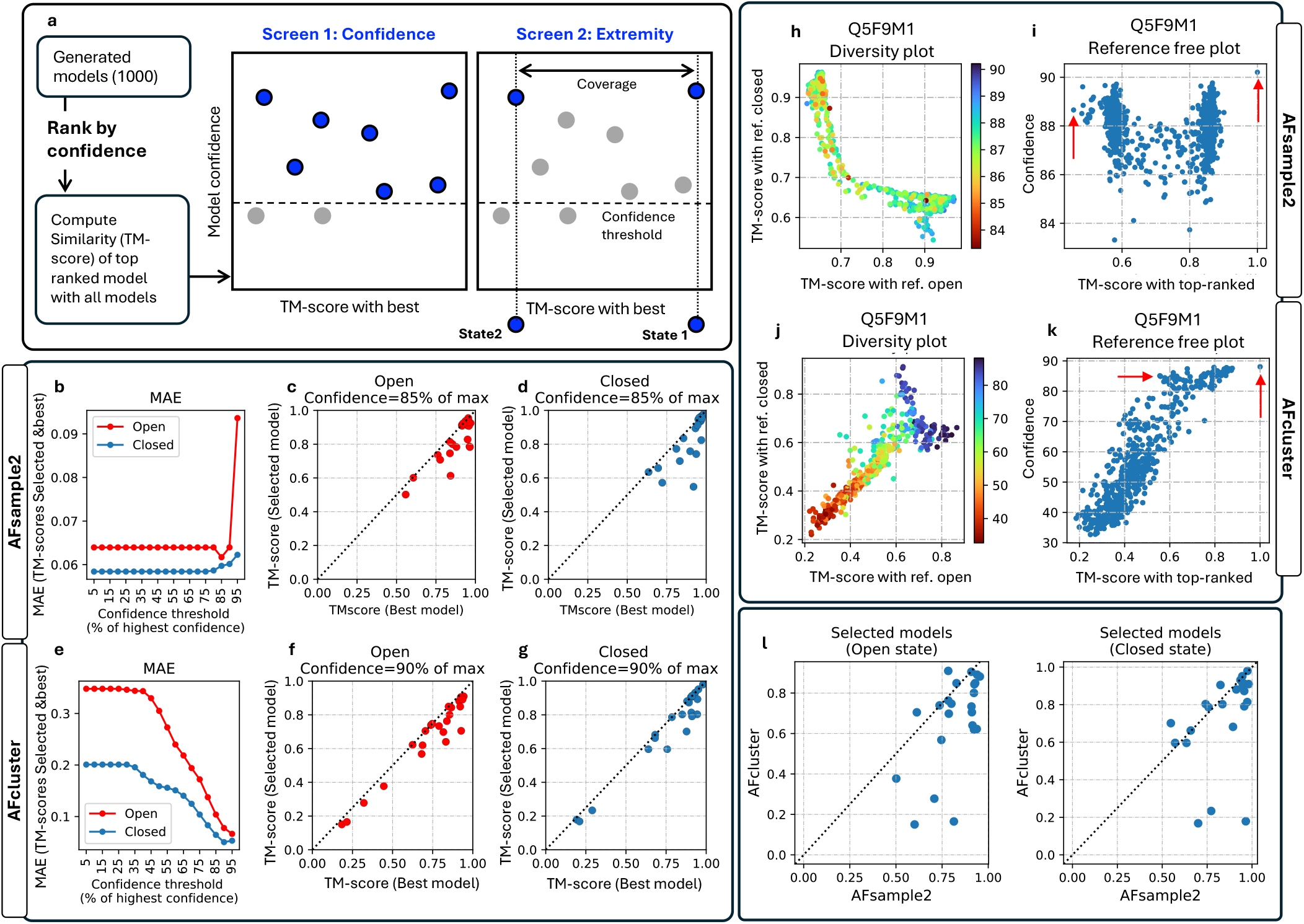
Reference-free state determination. (a) Schematic illustrating the process of identifying open and closed states from a pool of generated models. (b, e) Mean squared error (MSE) between the TM-scores of selected and best possible model for AFsample2 and AFcluster. (c, d) Scatter plot between TM-scores of selected models and best possible models generated by AFsample2 for all targets in the OC23 dataset (f,g - for AFcluster). An example with target Q5F9M1 showing (h) diversity plot and (i) reference-free plot. The reference-free plot has been annotated with optimal selections for the open and closed state (j, k - AFcluster). (l) Comparing selected open (left) and closed (right) models with the reference-free strategy for AFsample and AFcluster.

The strategy to identify the alternate state was bench-marked on model ensembles generated by both AFsample2 and AFcluster individually to ascertain the robustness of the method across varied ensemble characteristics. The quality of the selected states was assessed using the mean squared error (MSE) between the TM-scores of the selected states and the best possible selection (determined using reference structures). The *confidence screening*, step (ii), requires a threshold, and we monitored the MSE as a function of this threshold for AFsample2, and AFcluster, Fig 6b,e. We choose to make the threshold relative to the highest confidence rather than absolute to account for predictions of varying quality. For AFsample2 there is actually almost no need for a threshold, as the selection is equally good even without any threshold (0% of highest). For AFcluster, on the other hand, a threshold is needed since those ensembles contain many low-confident models, exemplified in Fig 6j. Around 90% and 85% of the highest confidence seems to be the optimal threshold for AF-cluster and AFsample2, respectively. But ideally, it is best to manually inspect the reference-free plots Fig 6i,k to locate the alternative states. For the optimal threshold the MAE is around 0.06 TM-score units for both AFsample2 and AFcluster, however, the MAEs are not comparable since the ensembles are different.

The actual similarity of the selected states and the best possible selection is summarized in Fig 6c,d, and f,g for AFsample2 and AFcluster, respectively. In almost all cases, the selection is relatively close to the optimal selection. AFsample2 has one poor selection (MSE*>*0.27) of the closed state for target B7IE18, the selected models is 0.55, while the best is 0.92. The selected models by AFcluster have no such failures, but since the AFcluster ensembles are of lower quality, the risk of failure is also lower. Reference-free plots for all targets in the OC23 dataset are shown in Fig S6

The selected states for target Q5F9M1 are illustrated in Fig 6h,i and j,k for AFsample2 and AFcluster, respectively. Despite the difference in the model ensembles between AFsample2 and AFcluster, the proposed method to identify the alternate state is able to select relatively high-quality models of both states irrespective of the method used for generating the models. This is true for most targets in the OC23 dataset, see Fig 6l.

### Evaluating AFsample2 on additional datasets

As described earlier, the AFsample2 pipeline performs well in predicting both the open and closed state on the OC23 dataset. However, it is important to demonstrate the same level of effectiveness for additional datasets. To this end, a dataset of 16 transporter proteins, which contains the ‘inward facing’ and ‘outward facing’ conformational states, was utilized (see Methods). 1000 models were generated for each method and the result was analyzed. We first compared the best generated models between AFsample2 and AFvanilla, see Fig. 7a. It should be noted that this dataset is fundamentally different from the OC23 as models generated by AFvanilla do not show a bias towards one of the conformations. Regardless, AFsample2 was able to generate superior models for both states, as illustrated by the improvement in ΔBest TMscore compared to AFvanilla for almost all cases 12/16 the alternate state is improved without deteriorating the performance of the AFvanilla preferred state. In two cases, the inward-facing is improved on the expense of the outward-facing, and in two cases there is no improvement at all. The best generated models for each target are shown in Fig. 7b, for AFvanilla, AFdropout, AFcluster, and AFsample2, respectively. Here, the models generated by AFsample2 cluster in a tight distribution at the top-right of the scatter plot. This improvement can also be illustrated in the plot between the fraction of successful proteins, defined as having both states above a certain TM-score (TM-score>threshold), and threshold, see Fig. 7b. At threshold TM-score>0.6, AFsample2 is successful for 90% of the targets, while the other methods only succeed for 50% of the targets.For most targets, AFsample2 generates the best models for both the outward-facing and the inward-facing states, see Fig. 7d,e. Only for two targets for the outward-facing and one target for the inward-facing does AFcluster generate the best models.

**Fig. 7.**
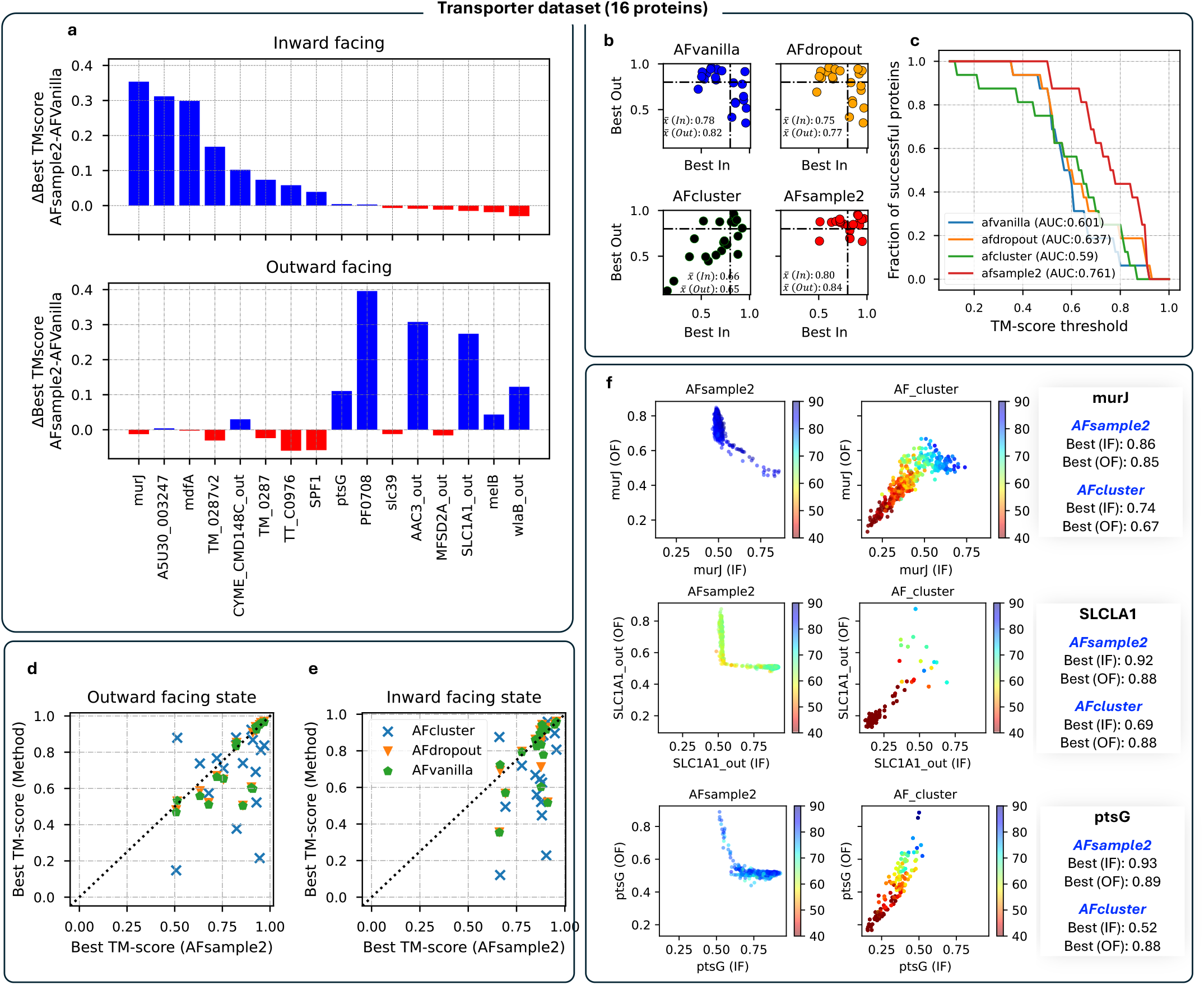
AFsample2 tested on a transporter dataset with 16 targets. (a) Improvement in best inward and outward facing conformations when compared with AFvanilla.(b) Best generated end states among methods for the transporter dataset shows the effectiveness of AFsample2. (c) Additionally, the success rate at multiple TM-score thresholds is summarized for all methods under consideration. (d,e) Best generated model for inward-facing and outward-facing states for all 16 targets showing that AFsample2 generated better models in most cases. (f) A subset of targets from the transporter datasets that could not be modeled in the original study (17). The diversity plots clearly indicate the effectiveness of AFsample2 in generating high-quality models of both states.

Furthermore, the original work (17) that constructed the transporter data set attempted to model the inward- and outward-facing states with multiple methods. While many methods succeeded for some cases targets, none of the methods were able to generate alternate conformations for these targets: *murJ, SLCLA1*, and *ptaG*. Fig. 7g summarizes the results when these targets were run with AFsample2. It can be clearly seen that AFsample2 is able to generate both states with a reasonable level of confidence, while AFcluster fails to generate the alternate conformation.

Apart from the presented comparisons, another interesting observation with regard to the confidence of models generated by AFcluster. It can be observed from Fig. 7f and Fig. S7 that a substantial number of AFcluster models have a very low confidence. This is the result of some of the clustered MSAs lacking information to generate high-confidence models. On the other hand, randomized column masking in AFsample2 is able to induce diversity without removing information that is crucial for generating high-confident models.

### Case study: Modelling un-modellable fold-switches with AFsample2

The inability of AF2, even with MSA sub-sampling, MSA clustering, and shallow MSA, to model intrinsically disordered proteins/regions (IDP) and fold-switch proteins has been reported (18, 19). This limitation has largely been attributed to the lack of IDPs in the training set of AF2. Although methods like AFcluster were originally developed and tested on fold-switch proteins, the results in a recent study indicate the ineffectiveness of all MSA sampling methods in modeling a modified S6-ribosomal protein that alters conformations between a *α*/*β*-plait (FS1) and a 3*α*- helical (FS2) fold (20) dependent on temperature, see Fig 8.

**Fig. 8.**
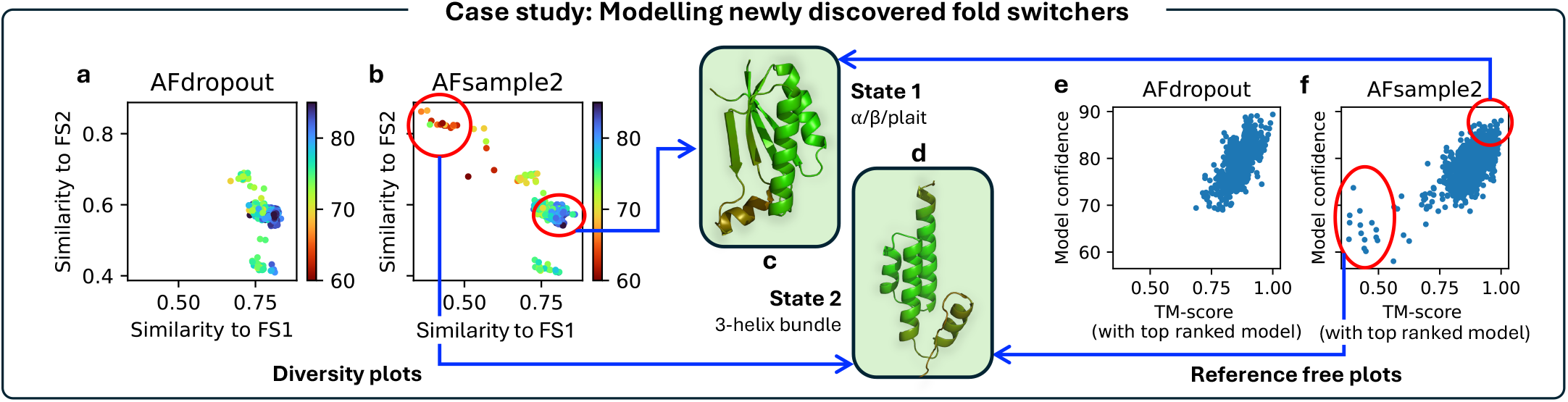
Modeling fold switchers with example of a S6-ribosomal protein. (a, b) Diversity plot showing the similarity of the generated ensemble to both folds. The models associated with FS1 and FS2 are annotated with (c, d) structures of both folds. Furthermore, the alternate fold can also be visualized on the scatter-plot generated with the (e, f) reference-free state determination protocol for the AFsample2 ensemble.

We generated 1000 models with AFsample2 and AFdropout for the modified S6-ribosomal protein, indeed the majority of the generated high-confident models are similar to the FS1 state (TM-score*>*0.8), this is true for both AF-sample2 and AFdropout. However, for AFsample2, there is a handful of models that are highly similar (TM-score*>*0.8) to the FS2 state, see Fig 8. These models have lower model confidence compared to the FS1 state (0.75 vs 0.88), but from the reference-free plot, they can still be identified as a relatively confident alternate state. Generating more models could potentially yield higher-scoring models for the alternate state.

Current efforts that use AF2 for modeling alternative states of disordered proteins rely on model confidence to select models. However, as AF2 was trained to estimate high quality models for a single conformation, the confidence metric might not be a reliable indicator while looking for alternative states as the AF2 inference system is bound to rank these structures lower. This was also observed in the OC23 set, where the open conformation was consistently less favorable in terms of model confidence Fig S2. Instead, above a minimum confidence score, the similarity to highest confident state would serve as a better indicator of model diversity.

In summary, current methods that aim to improve conformational diversity tend to be sub-optimal for fold-switch proteins. AFsample2 shows that it is possible to utilize the AF2 inference to make predictions for proteins that have not been part of its training set and that AF2 can generalize to new unseen proteins.

## Discussion

Sampling and clever MSA reshuffling have been shown to improve the conformational coverage of models generated using the AF2 inference system. However, solutions are required to estimate better (i) end-states as well as (ii) intermediate states to understand the full picture of the conformational dynamics. Additionally, there are many unsolved questions, such as how to correctly identify states without a priori knowing the structure of the state and how the states are physically and dynamically connected. In addition, we are struggling with insufficient experimental data where, in many cases, the structure of only one state is known, usually the most stable under experimental conditions, and we cannot tell with certainty if our predictions are correct. Of course, this is also an opportunity for computational methods to provide experimental hypotheses that can be used to validate the predictions.

A major strength of the AF2 inference system is its efficient extraction of the co-evolutionary structure embedded in MSAs. This capability is highly beneficial for generating a single, high-confidence model. However, when attempting to expand the conformational landscape of predictions. This specific feature of the AF2 system becomes a drawback, as the strong co-evolutionary constraints prevent the generation of alternative states.

As a solution to these problems, we developed AFsample2, a method to generate high-quality conformational ensembles for proteins by randomized MSA column masking and a novel approach to identifying conformational states without the aid of reference experimental structures. We introduce noise within the MSA feature profile that leads to a significant improvement in the estimation of alternative states as demonstrated on diverse sets of proteins. Under the assumption that MSAs contain information about the patterns of correlated mutations (co-evolution) between residues that can inform physical contacts within the protein structure, making changes to these relationships is a viable strategy for inducing diversity in predictions. The idea is not limited to the AF2 inference system but could be readily applied to other MSA-based prediction systems, such as RosettaFold or OpenFold.

In addition, we used AFsample2 here to predict different states, but it could also be utilized to improve the conformational sampling of difficult single-state proteins that seem to be stuck in one local minimum. In terms of limitations and future directions, the current version of AFsample2 has only been tested on monomers. However, it can be easily adapted for generating conformational states for multimeric protein complexes as well.

## Methods

### AlphaFold version

The entire prediction pipeline is based on AlphaFold v2.3.1, which is available at https://github.com/google-deepmind/alphafold. The original transformer model assumes that the embedding vectors do not correlate to each other (21). However, in the case of the AF2’s Evoformer, the relationships between MSA rows (alignments) and columns (residues) were established with row-wise and column-wise attention, respectively (8). Randomized MSA masking is aimed at diluting these coevolutionary signatures and increasing the probability of sampling alternative states.

### Datasets

Below, we describe the different datasets used in this study.

#### OC23

A subset of 23 proteins with distinct open and closed states were selected as defined by a TM-score difference between the open and closed state *<*0.85. An overall summary of this dataset with their PDBIDs is compiled in Table S1.Protein sequences extracted from the experimental open and closed structures could have been directly used to run AF-sample2 predictions. However, due to irregularities such as (i) Unequal chain length, (ii) missing residues, and (iii) in-complete information, we chose to retrieve the full protein sequence from Uniprot. With this, we were confident that the inference system was getting the complete information regarding the protein’s sequence. In summary, all predictions were performed on the full protein sequence from UniProt for each protein.

#### Transporters

For the validation sets, a transporter dataset with 21 unique proteins with inward- and outward-facing conformation was utilized. This dataset was manually cu-rated in a recent study to benchmark protein structure pre-diction methods for membrane proteins that show substantial conformational changes (17).

### MSA generation and randomization

The MSAs were produced using the DataPipline implemented in AF2with default parameters using HHblits (22) and Jackhammer searches on Uniref90 (23), BFD, Uniclust30 (24) and MG-nify sequence databases. Templates were not used in any of the runs. Randomized MSA masking was incorporated into the pipeline at the MSA pre-processing step. The algorithm sequentially initializes model runners associated with specific model names on a given FASTA sequence. Notably, the key innovation lies in the introduction of variability through MSA randomization, a process governed by the user-defined parameter ‘msa_rand_fraction’. When this fraction exceeds zero, a duplicate of the original feature dictionary is generated and the given fraction of MSA columns is randomly selected and replaced with ‘X’.

#### Inference

Since multiple versions of trained model weights are available for AF2, the choice of the weights was an important consideration. AF2 has two versions of the monomer neural network weights, with each version having an additional five sets of neural network weights. The first version used during CASP14 was extensively validated for structure prediction quality (8). The additional five pTM models were fine-tuned to produce pTM (predicted TM-score) and (PAE) predicted aligned error values alongside their structure pre-dictions. The relative difference in these models was assessed in terms of the TM-score profile of the predictions with experimental open and closed states. All ten model-types were tested individually to ascertain if any of them was consistently doing better than the others. The results summarized in Supplementary Fig S3a, show that every model-type has the ability to generate the best model depending on the protein. Also, Fig S3b indicates that the quality of models in terms of confidence did not vary much with different model weights. Based on this, and to achieve maximum performance, the AFsample2 pipeline utilizes all ten neural network weights for inference.

#### Diversity analysis

The proposed measure to quantify conformational diversity given a set of protein models was developed. It performs the following tasks. (i) *Point Projection and Transformation*: Project points onto a line defined by two endpoints *A* and *B* and followed by a 45-degree counterclockwise rotation. (ii) *Histogram Generation*: Calculation of a histogram of the transformed points and quantifying the fill ratio as the ratio of non-empty bins to the total number of bins in the histogram. (iii) *Data Visualization*: Plot histograms to visualize the spread of models in a plot of TM-score with open conformation vs TM-score with the closed conformation. A formal summary of the entire process and a visual description can be seen in Fig 9.

**Fig. 9.**
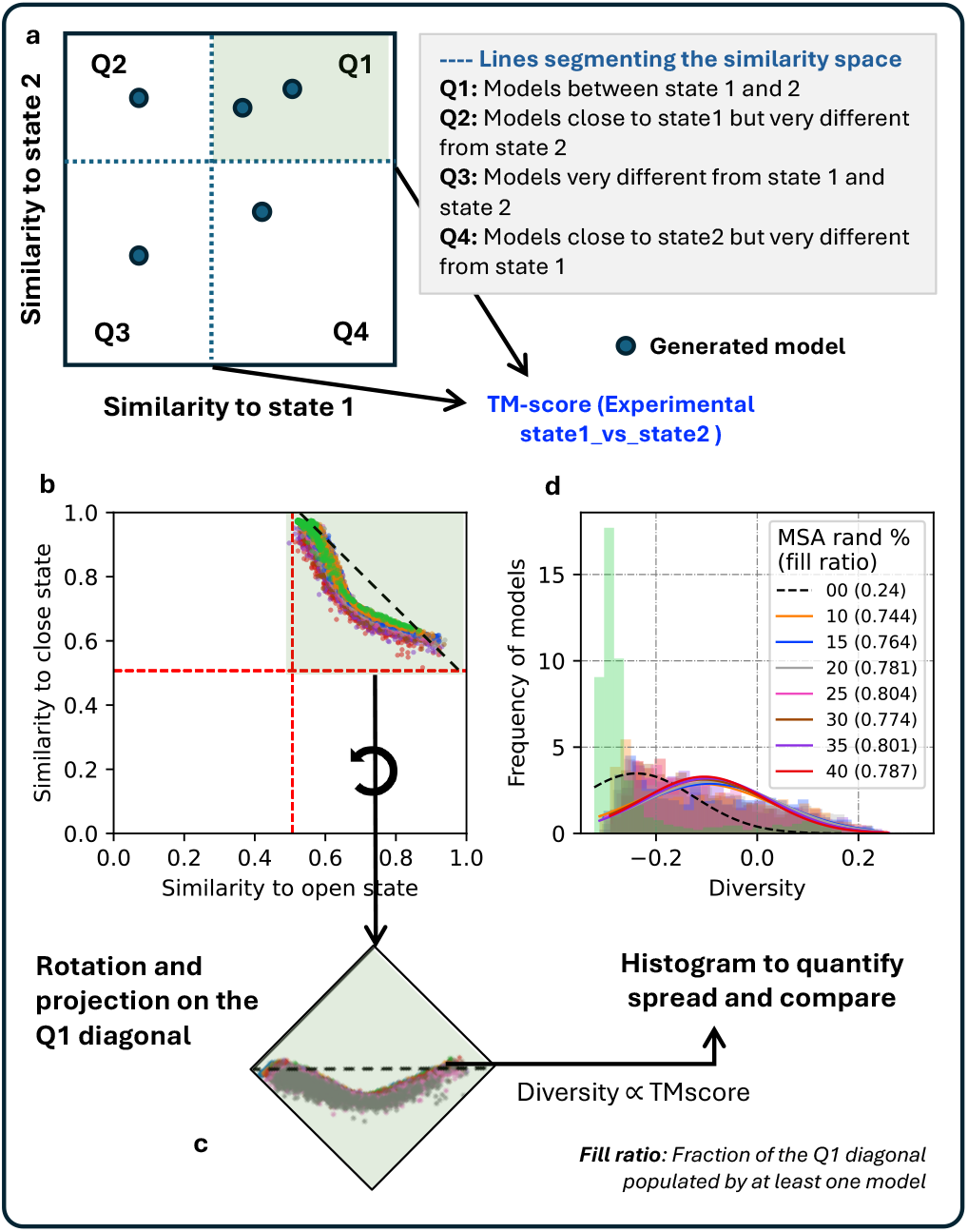
An overview of the fill-ratio metric. Given (a) diversity plot, this protocol quantifies the conformational diversity of ensemble models. (b, c) All models in Q1 are projected onto the diagonal. Models in other quadrants (Q2, Q3, and Q4) are ignored since they are not close to any of the two reference states. The diagonal is then divided into 100 bins and the ratio of bins containing at least one model is recorded. This ratio gives information about the conformational space spanned by the model ensemble between the two reference states.

Let *A* and *B* be the endpoints of the line onto which points are projected. These points are defined as (1, TMoc) and (TMoc, 1), where TMoc is the TM-score between the experimental open and closed conformation. Further, *P* is a point (denoting a protein model) that has to be projected onto the line. *AP* be the vector from *A* to *P* . *AB* be the vector from *A* to *B*. The projection of point *P* onto the line passing through *A* and *B* can be calculated as:

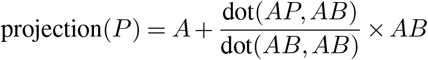

where dot(*x, y*) denotes the dot product between vectors *x* and *y*.

The rotation of the projected points by an angle 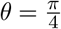 counterclockwise can be achieved using the rotation matrix:

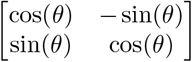

The histogram is generated with a specified number of bins, and the fill ratio is calculated as the ratio of non-empty bins to the total number of bins. This can be applied in the analysis of protein structures to (i) Analyze the spread of models in the TM-open vs TM-closed plot, (ii) Quantify the degree of overlap or clustering of data points, and assess the quality and completeness of data representation.

### Investigating the effect of sampling on conformational diversity

The impact of sampling size on the accuracy and reliability of model state estimations, particularly within biological datasets categorized by protein states (‘TM_open’ and ‘TM_closed’), was investigated. This analysis was aimed at quantifying how different sampling sizes influence the derived states’ quality and reliability. For each protein identified within a batch, the dataset was bifurcated into two subsets based on the state of the protein (either open or closed). A series of experiments were then conducted to ascertain the variability of the maximum TM-score (a metric indicative of model similarity to a particular state) with an increment in sample sizes, ranging from 50 to 1000 in steps of 50. In each experimental iteration, a specific number of instances were randomly selected (n) from the model ensemble, with the maximum TM-score to open and closed state recorded. This process was repeated 1000 times for each sampling size to reduce the bias associated with random selection, and the mean of these maximum scores was computed to represent the score for that particular sampling size. This method was applied across all proteins within each batch, compiling the mean maximum scores for both open and closed states. The results were aggregated to analyze the overall trend across all batches.

### Evaluating relative performance of difference randomization levels

The dataset comprises multiple instances of protein predictions, each associated with a unique identifier (UniProt ID) and categorized under different configuration settings (randomization %). For each unique protein, predictions were further characterized by their state (TM_open), and their similarity to the experimental open state was quantified using the TM-score. To systematically compare the performance of configuration settings, the Wilcoxon ranksum test, a non-parametric statistical test, was adopted to evaluate the hypothesis that one configuration yields higher TM-scores than another. Given the multiple comparisons issue inherent in our analysis—comparing each configuration against all others for each protein—a Bonferroni correction was applied to adjust the significance threshold, thereby controlling the family-wise error rate. Specifically, the original alpha level of 0.005 was divided by the total number of comparisons (36), setting a more stringent criterion for statistical significance.

### Evaluation metrics

TM-score (25) calculated using TM-align (26) was used as the evaluation metric for comparing similarity to reference states and to other models. TM-score was calculated using a fixed *d*_0_ = 3.5Å to avoid the problem of long proteins achieving artificially high TM-score (25).

## Data availability

All scripts and data presented in this study are made available at https://wallnerlab.org/AFsample2

## ACKNOWLEDGEMENTS

This work was supported by the Wallenberg AI, Autonomous System and Software Program (WASP) from Knut and Alice Wallenberg Foundation (KAW), Swedish Research Council grant, 2020-03352, The Swedish e-Science Research Center, and the Wenner-Gren Foundation. The computations were performed on resources provided by KAW and NSC (Berzelius).

## Supplementary Figures and Tables

**Fig. S1.**
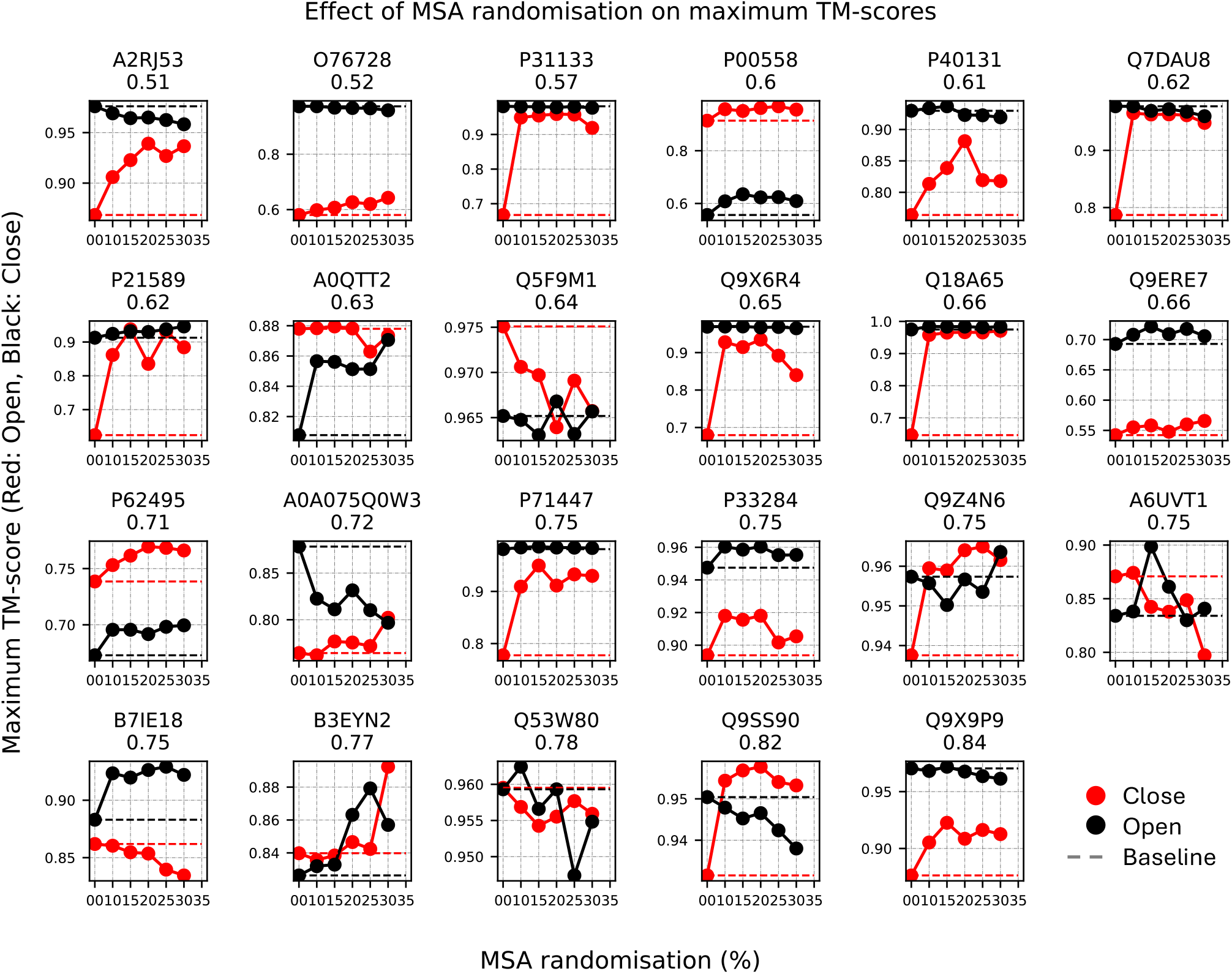
Best selected models by AFsample2 at different levels of randomized MSA masking in the OC23 dataset.

**Fig. S2.**
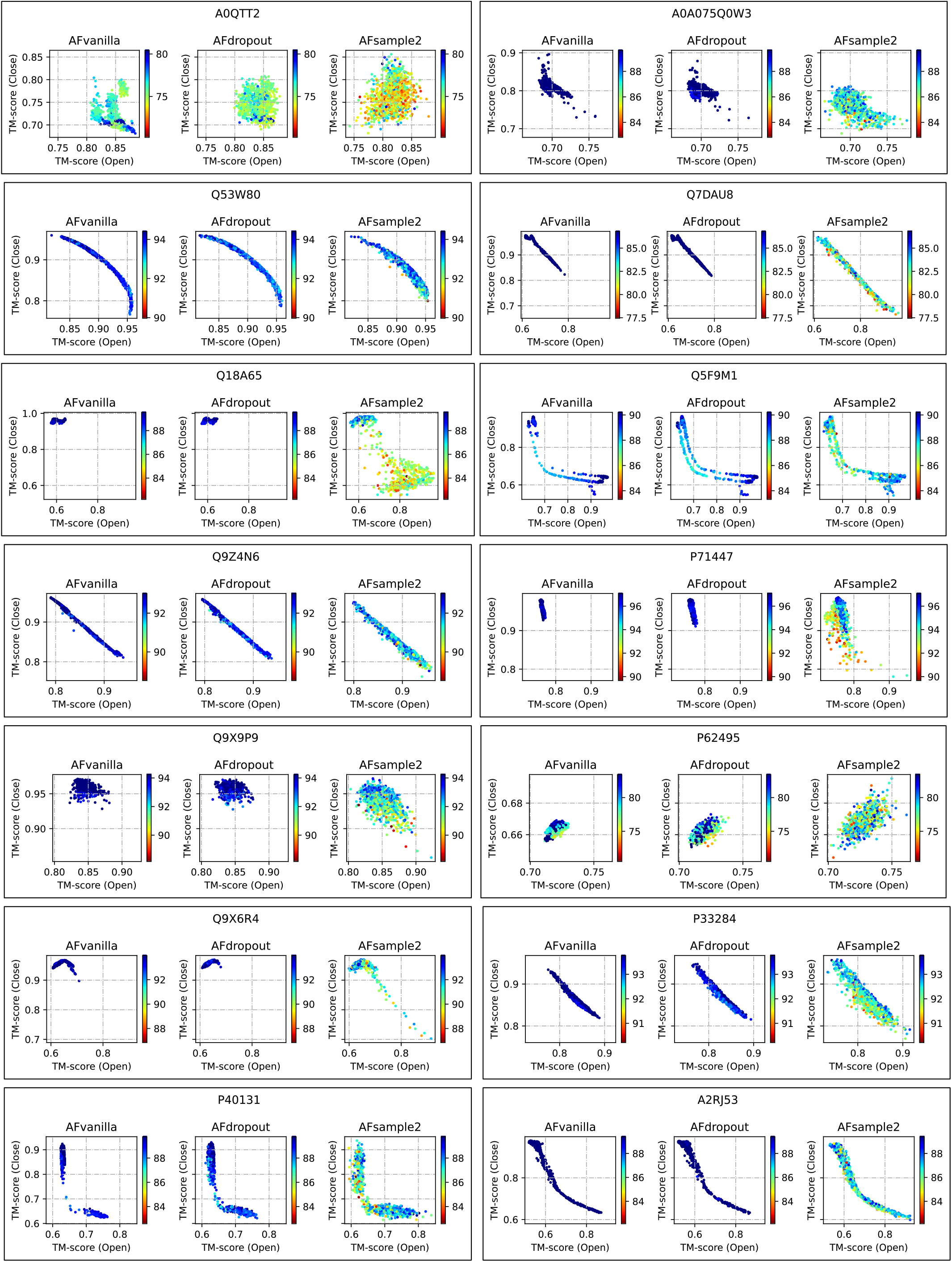
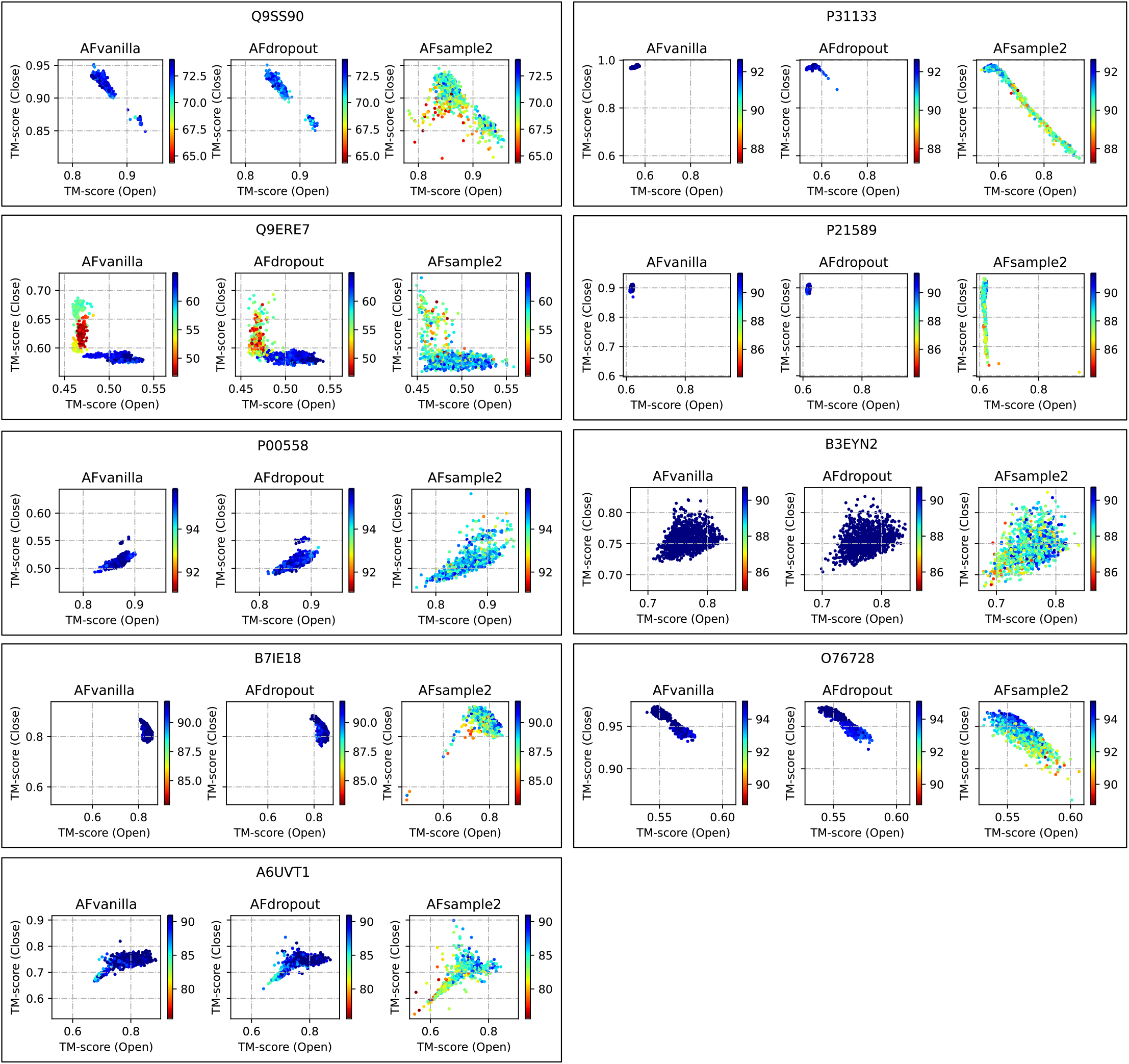
Diversity plots to capture the similarity of model ensembles with reference states (continued to next page)

**Fig. S3.**
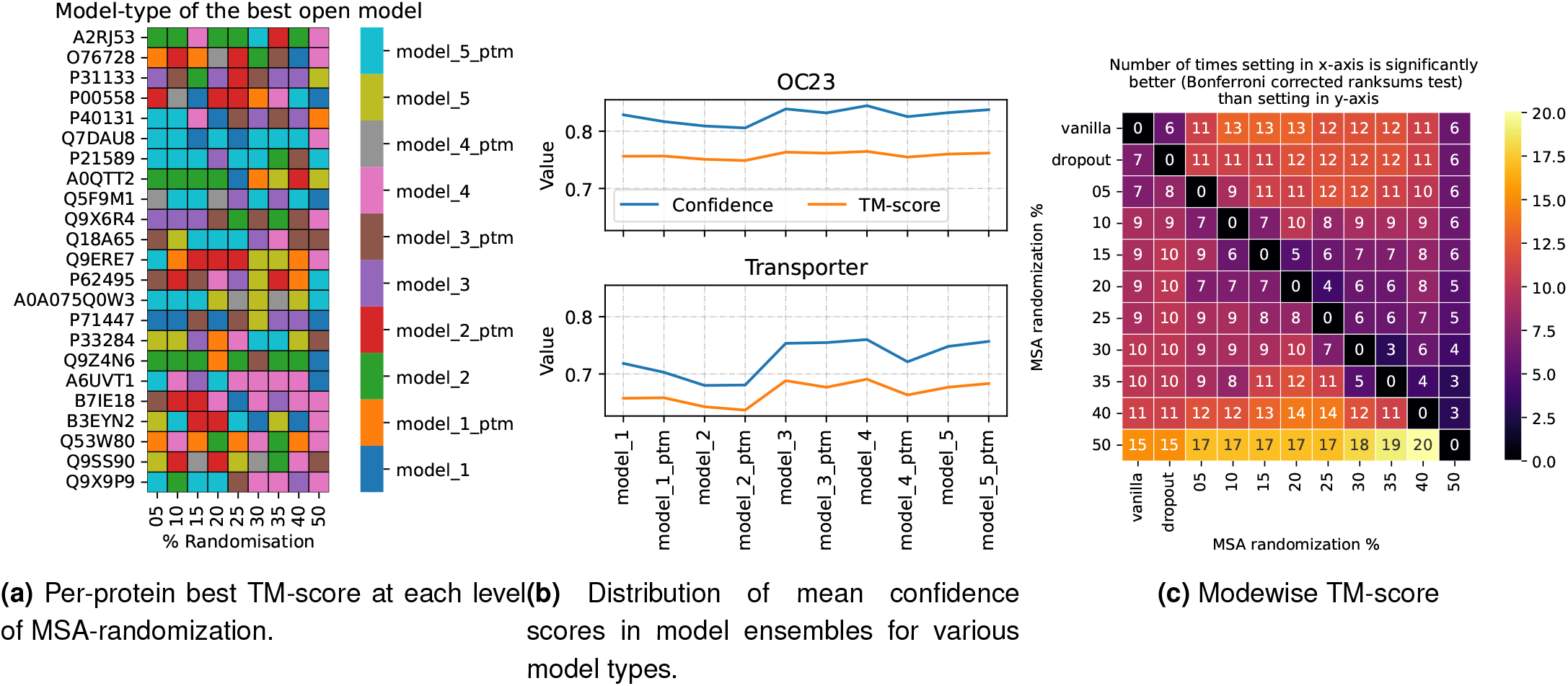
(a) Analysis to see if any of the model-types are favoured to generate the best-open model. Since the heatmap does not show any clear pattern, it can be inferred that model-type preferences is completely dependent on individual protein.

**Fig. S4.**
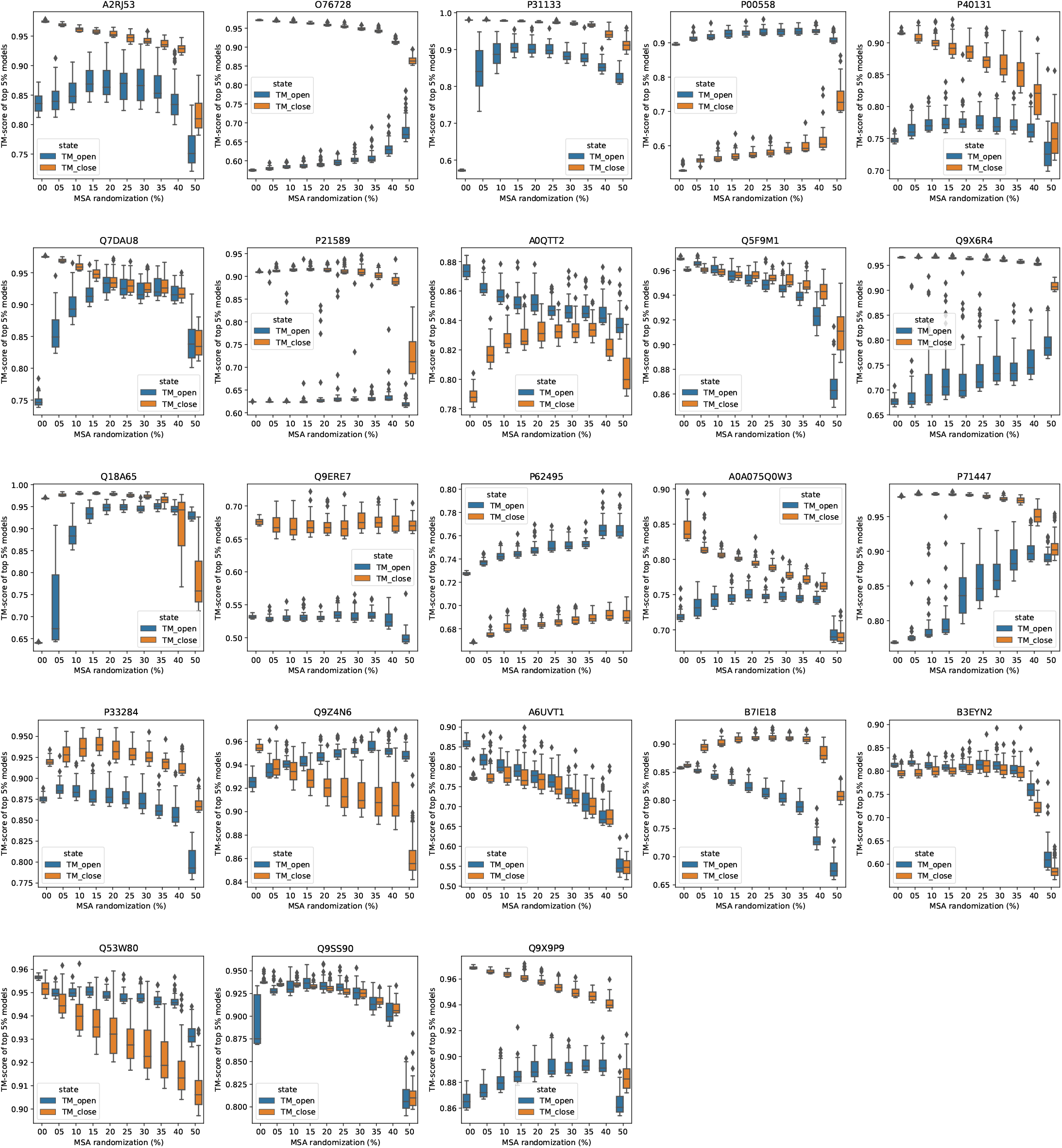
Changes in TM-scores for the top 5% open and closed models for all proteins with MSA randomization in the OC23 dataset

**Fig. S5.**
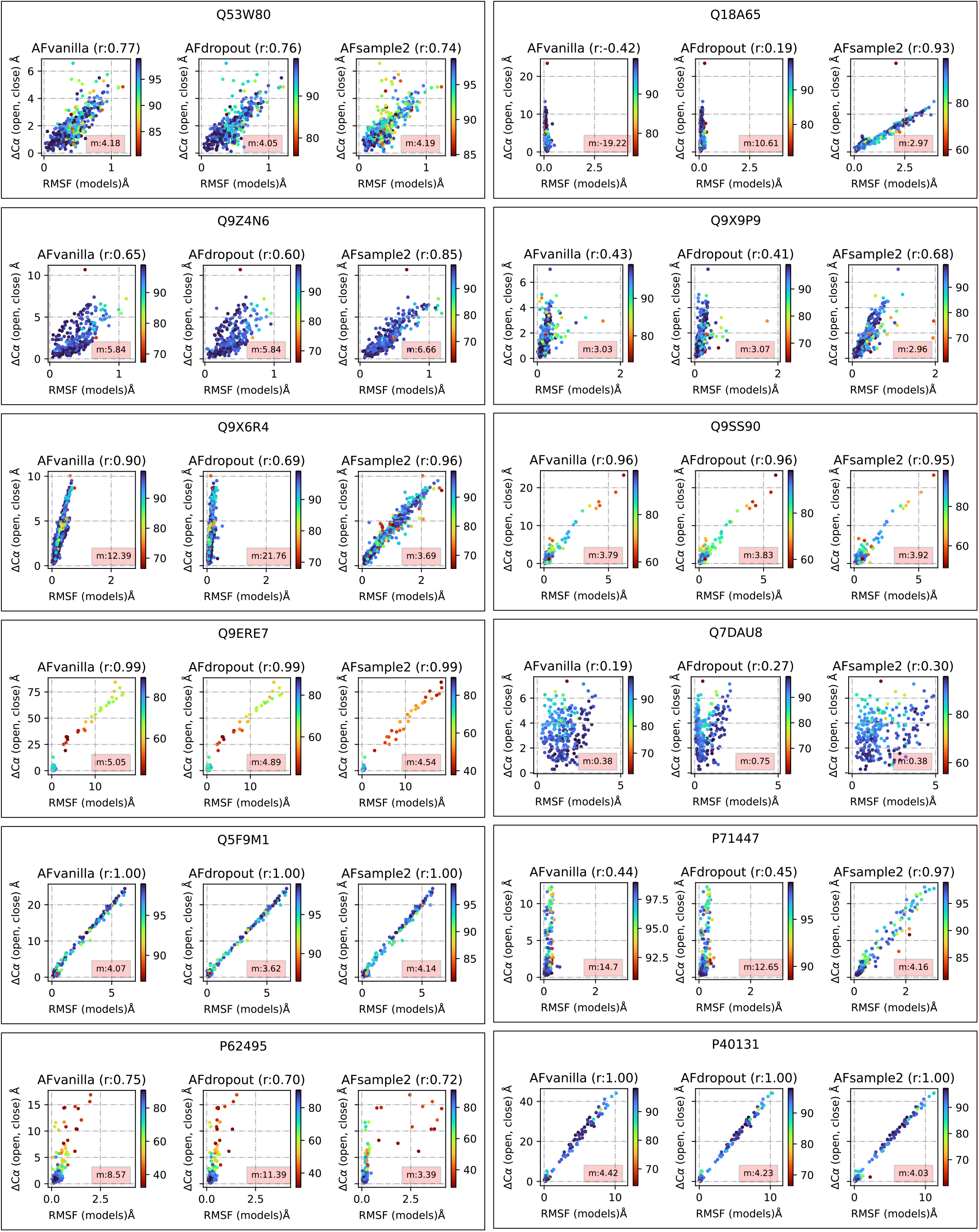
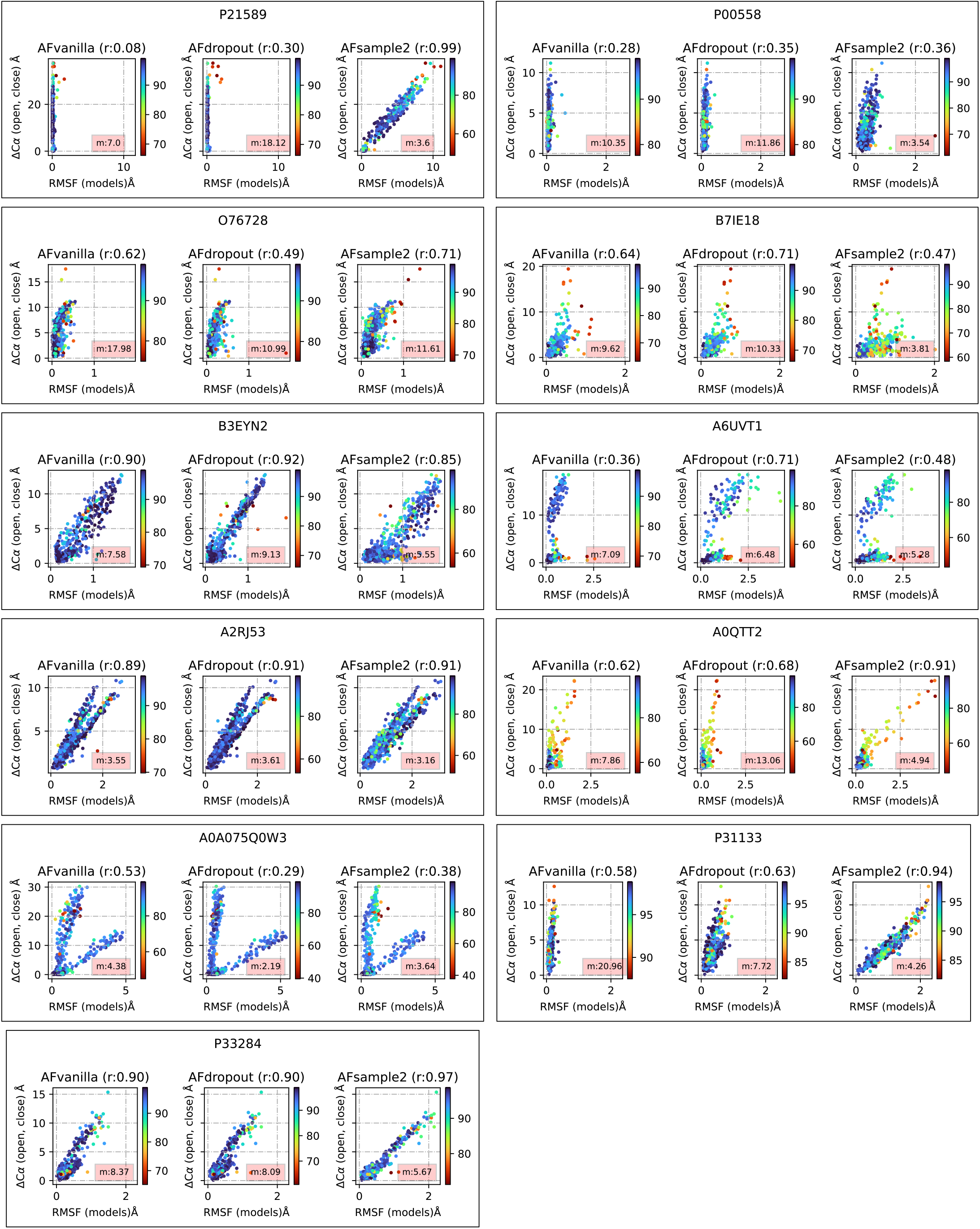
Correlation between fluctuations observed in the generated model ensemble and reference states at a residue level (continued to next page)

**Fig. S6.**
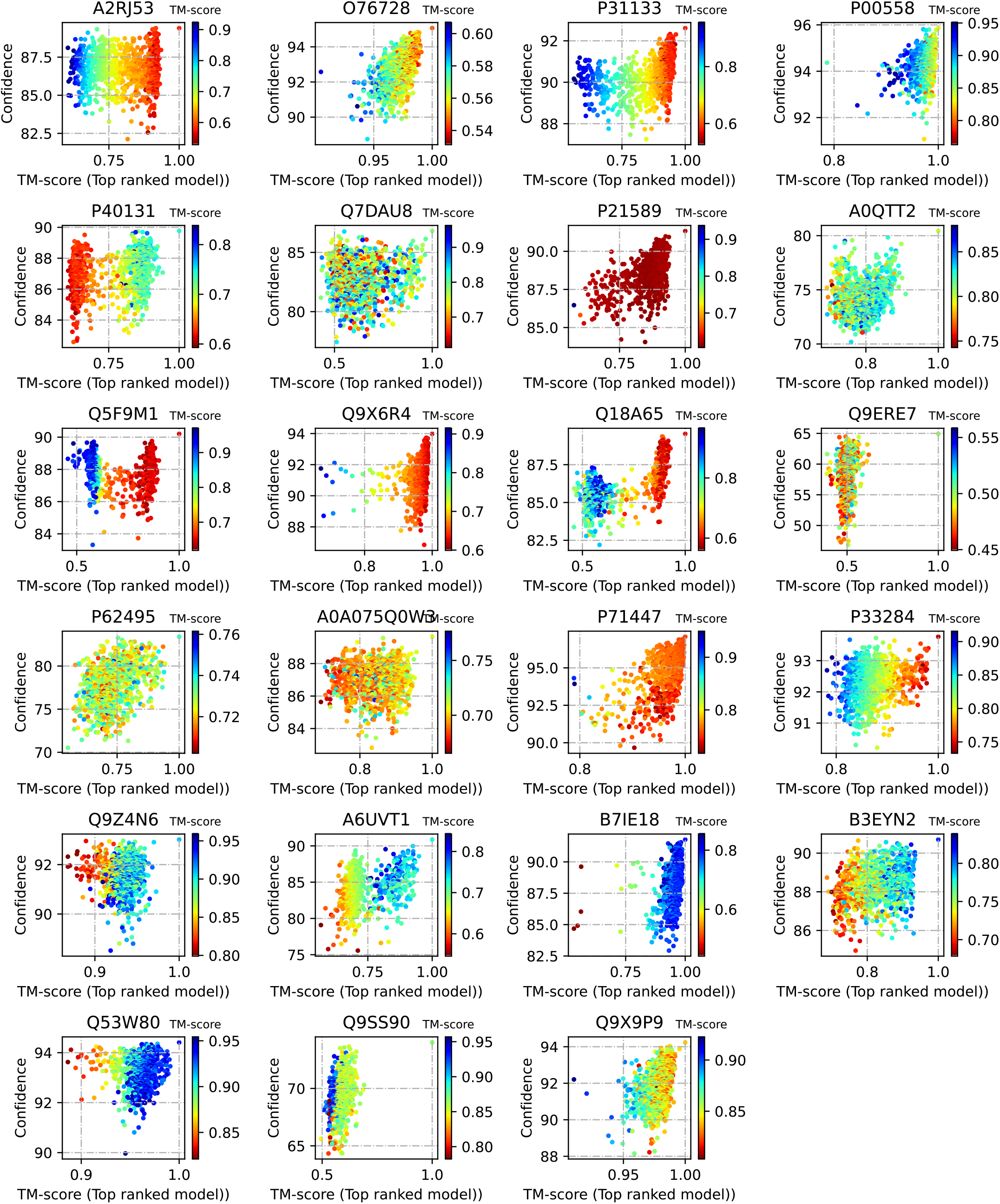
Reference-free state determination. The assumption regarding alternate state to be far from the most confident model holds for the OC23 dataset. The color-gradient denotes closeness to the alternate (open) state.

**Fig. S7.**
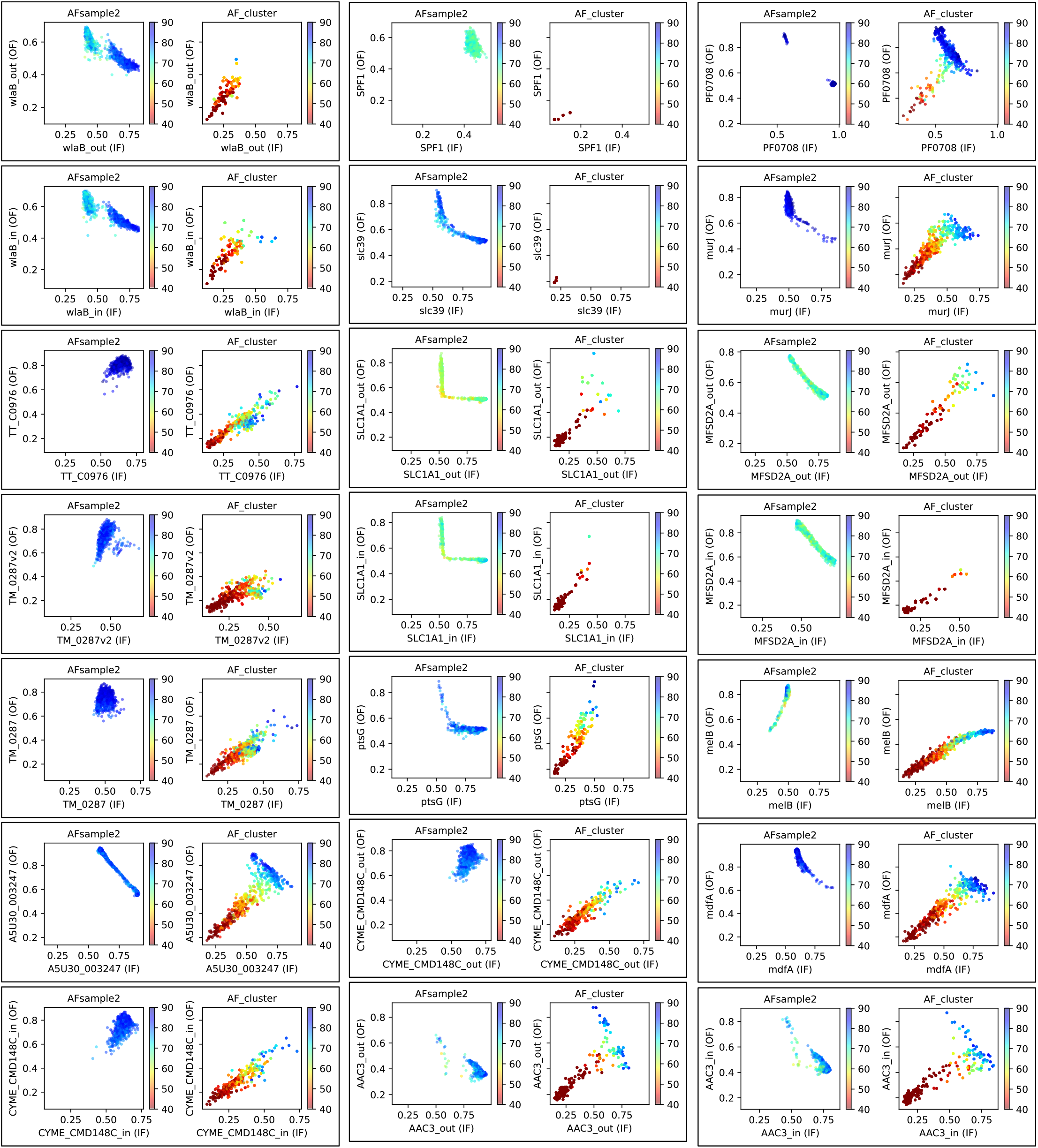
Comparing diversity plots for all 21 proteins in the transporter dataset for AFsample2 and AFcluster.

